# Establishing Normative Reference Values for the Utah Seated Medicine Ball Throw Protocol in Adolescents

**DOI:** 10.1101/2021.12.14.472637

**Authors:** Cory Biggar, Abigail Larson, Mark DeBeliso

## Abstract

The seated medicine ball throw (SMBT) is a field test intended to assess upper-body muscular power by measuring the maximal distance an individual can throw a medicine ball from an isolated, seated position (25). The SMBT has been used to assess upper-body power in various populations and to establish concurrent validity for other measures of upper-body power such as the bench press power test and the plyometric push-up. The SMBT is less costly and simpler to incorporate into a field test battery than other upper body power assessments. While the SMBT is a valid, reliable field test for upper-body power, normative reference standards for most populations, including adolescent (12-15 years old) physical education students, do not exist.

**Purpose:** This study reports distances thrown in the SMBT to establish normative reference values in male and female physical education students, ages 12-15 years old.

**Methods:** One hundred thirteen untrained male and female physical education students aged 12-15 years performed the SMBT field test three times on a single testing day. Participants threw a 2kg medicine ball with a 19.5 cm diameter while seated at a 90° after recording height, body mass, and BMI.

**Results:** Participant data was separated by age gender for analysis. Mean and standard deviation for the SMBT for males was 4.3±0.7m and 5.2±0.8 m for ages 12-13 and 14-15, respectively, and for females was 3.4±0.5m and 3.7±0.5m for ages 12-13 and 14-15, respectively. Pearson correlation coefficients for between-trials comparisons for males and females ranged from r=0.85-0.97. Pearson correlation coefficients for average SMBT and age of participants was r=0.93. Normative reference values as percentile ranks for the SMBT scores for age groups 12-13 and 14-15 among males and females, respectively, were also established.

**Conclusion:** The data presented provides an initial set of normative reference standards for coaches and students to determine upper-body muscular power using the SMBT.

## INTRODUCTION

Muscular power is an essential athletic performance variable within many sports and has been researched extensively (2). Beachle and Earle highlight the importance of power, describing it as the attribute that allows athletes to overcome gravity, accelerate the body through space, and accelerate a ball across the court or playing field (2). Power, in most cases, refers to a high rate of coordinated, forceful contraction of the muscles, controlled by numerous factors, including type muscle action, mass lifted, the architecture of muscles, fiber cross-sectional area, range of motion, and movement distance (26). Mathematically, power is work divided by the elapsed time when work is equal to force multiplied by the movement distance (2).

Due to the nature of the skills and techniques required, upper-body muscular power plays an especially significant role in sports such as basketball, cheerleading, volleyball, tennis, and gymnastics (4, 16, 27). Whether it is the athlete’s body or a foreign object such as a ball, the ability to accelerate objects through space is essential for many sports. When assessing readiness/aptitude for sport, muscular power is a vital consideration. Part of the task for physical educators is to prepare students for a lifetime of physical activity through sport and lifetime activities. Therefore, it is prudent for physical educators to assess and track upper-body muscular power to assess the success of the physical education curriculum and prepare students for sport participation.

As a construct, upper-body muscular power should be easily measurable and comparable to normative reference values. While many methods currently exist for measuring upper-body muscular power, convenience, cost, prerequisite physical requirements and feasibility vary across testing protocols (5, 10, 17, 30). Many upper-body power assessments, such as the bench press power test, are technique-intensive and require specialized equipment, thereby limiting their practicality when aiming to assess larger groups of non-resistance trained individuals. Alternately, the seated medicine ball throw test (SMBT) is a field test that assesses upper-body muscular power, specifically in the pectoralis, shoulder, and elbow flexor muscle groups, and represents a practical and safe, reliable testing method. The SMBT assessment requires an individual to throw a medicine ball from an isolated, seated position, the test administrator then measures the distance thrown from the start position to the first contact point (5). The SMBT is less costly and less complicated to incorporate into a testing battery than other assessments such as the bench press, rope-climb, pull-up, and force-plate plyometric push-up as it requires little technical or equipment expertise and minimal prerequisite strength and technique requirements (6, 8, 10, 30).

The SMBT is a valid and reliable measure of upper-body power in various populations (table 1). The SMBT has a low coefficient of variation (CV) and high intraclass correlation coefficient (ICC) when examining variables such as maximum velocity (3.2 & 0.93 for CV and ICC, respectively) and acceleration (3.3 & 0.85 for CV and ICC, respectively) (29). In many cases, the SMBT test has been used to establish concurrent validity for other measures of upper-body power. Clemons et al. used scores from the SMBT to assess the validity of the bench press power test (6). The results from the study showed concurrent validity between the bench press power test and the SMBT (r=0.86, p < 0.01) (6). The SMBT is also strongly correlated to other tests of muscular power, such as the rope-climbing test (r=0.99, p<.05) and the Wingate test (r=0.655, p<.05) (10, 22).

**Table 1.** Previous research using the SMBT as a measure of upper-body power

| **Population** | **Age**<br>**(Years)** | **Results** |
| --- | --- | --- |
| Untrained Women<br>(24) | 55-64 | Upper-body power correlates with average distance ($p<0.05$, $r=0.606$) and maximum distance thrown ($p<0.05$, $r=0.552$). |
| Female Netball<br>Players (7) | 21.6±2.1 | Peak power ($r=0.913$) and mean power ($r=0.872$) were highly correlated with chest pass (med ball throw) distance. |
| Highly Trained<br>Male and Female<br>Overhead<br>Athletes (4) | 18-50 | Showed excellent test-retest reliability ($r=0.958$) in the SMBT. |
| Male and Female<br>Track and Field<br>Athletes (11) | 20.0 ± 0.5 | SMBT showed a significant ($p\leq 0.05$) connection to upper-body power. |
| Antiterrorism<br>Soldiers (10) | 24.1 ± 1.8 | Showed significant correlation between SMBT and other upper-body power tests ($p<0.05$ $r=0.55-0.98$). |
| Resistance-trained<br>men (20) | 21.5 ± 2.3 | Showed a 90% confidence limit to indicate the effect of whole-body vibration on performance. |
| Adolescent Male<br>Basketball Players<br>(1) | | Used to show significant ($p=0.000$) difference in throwing distance between different age groups. |
| Male and Female<br>Physical<br>Education<br>Students (13) | 15-16 | SMBT used to show a significant ($p<0.05$) difference in performance between a trained and untrained group. |
| Male and Female<br>Special Olympics<br>Athletes (22) | | Showed a strong correlation for mean power ($r=0.642$) and peak power ($r=0.659$) between Wingate test and SMBT. |

Note: Parenthesis in column 1 refer to reference number

Although the SMBT is a reliable field test for upper-body power, there are few normative reference values, which may explain why it is not widely incorporated into sport and physical education assessments. While there is data on the SMBT in older adults and kindergarten-age children, relatively little data has been collected in adolescents (5, 7, 8, 14, 19). The relative underuse of the SMBT has resulted in a lack of comparable normative reference values. Published normative reference values provide a baseline measurement by which practitioners can compare results and would likely increase the utilization of the SMBT as a means to assess upper-body muscular power. To the best of our knowledge, no normative reference values for the SMBT have been established for adolescent (12-15 years) physical education students.

In addition to the lack of normative reference values, there is no official testing protocol for the SMBT. Some studies use protocols requiring participants to sit at a 45° on a bench (6, 10, 11, 20), while others require a 90° angle against a wall (4, 13, 24, 29). However, both appear to be reliable measures, and throwing distances appear to be similar (4, 11, 26). For example, college-age men (age 20.3 ± 1.1) years) seated at a 45° threw the ball a mean distance of 4.1 ± 0.5 m, while a similar group (age 23.1 ± 3 years) seated at a 90° threw the ball a mean distance of 4.1±0.5 m (4, 11). The results of these studies indicate that throwing distances between participants seated at different angles are similar. The mass of the medicine balls used also varies across studies. Medicine balls ranging from 2kg to 9kg have been used (6, 10,11, 20, 29). Obviously, the use of a lighter ball allows for further throw distance. Typically, the mass selected for an assessment of upper-body power is dependent on a percentage of the participant’s 1RM bench press weight, however determining the 1RM requires substantially more time, prerequisite strength and technique, and additional equipment and personal resources (6, 10, 11, 20, 29). As such, comparing results across studies is difficult (11, 20). Normative reference values and a standardized protocol for the SMBT, including weight thrown, for all populations will provide context for scores and delimit past and future research findings.

Considering the aforementioned limitations, the purpose of this study was to develop a protocol and normative reference value data set for the SMBT for middle-school-aged (12-15 years) physical education students. The present study will help to provide another valuable tool for coaches and physical educators to use in assessing upper-body muscular power. The present study results will allow for the development of a standard to assess physical education students’ upper-body muscular power using the SMBT.

## METHODS

### Participants

A convenience sample of 113 male and female physical education students, aged 12-15 years, from northern Utah participated in this study. Researchers recruited individuals from physical education classes in a single public school in northern Utah. Participants were considered untrained.

Researchers obtained human subject approval by the IRB (SUU IRB Approval #24-032020b). Physical education teachers issued a public announcement to their classes and asked those who wished to participate in the study to obtain written parental permission and return the signed informed assent before or on the day of data collection. Researchers required participants to be between 12 and 15 years of age and free of injury or disease for inclusion in the study. Informed consent/parental assent was obtained from the participant and parent(s) prior to any data collection. Participation was voluntary, and participants were able to withdraw at any time without penalty. Before the testing protocol, researchers discussed procedures, possible risks or discomforts, benefits, and confidentiality of information with the volunteers. All personally identifiable information about participants was confidential.

### Instruments and Apparatus

For the SMBT, a 2 kg medicine ball with a 19.5 cm diameter was used, along with a measuring tape and gymnastic chalk. The medicine ball was a rubber “Champion Sports” brand ball and was 19.5 cm in diameter (image 1). The measuring tape (20 meters) measured distance increments in meters. Researchers used a Detecto 437 eye-level physician’s scale to collect participants’ body mass, measured in kilograms. A separate measuring tape was used to measure participant height, measured in centimeters.

### Procedures

Participants completed all testing within the gym of their regular physical education class on a single day. Researchers spent an additional school day giving information to potential participants and handing out informed assent packets. In total, the study required two days to recruit participants and collect data.

Data collection for this study occurred during the COVID-19 pandemic. Due to the pandemic, researchers took additional measures to ensure the safety of participants and researchers. In order to protect both researchers and participants from possibly contracting the virus, commonly touched surfaces, such as the medicine ball, were sanitized between every use. All participants were required to wear masks during the data collection, and participants were kept six feet apart at all times.

#### Warmup Protocol

On the day of testing, the researcher read instructions to students and demonstrated the assessment. Before participating in the SMBT on the day of testing, participants completed a brief questionnaire then were measured for height and body mass. The questionnaire asked the age and gender of the participant. After recording height, weight, gender, and age, volunteers participated in a warmup protocol. Directed by the researcher, the warmup protocol consisted of multidirectional shoulder movements similar to those used in the study by Borms and Cools (4). In total, the warmup protocol was two minutes in length and required the participants to jog in place for 30 seconds, perform thirty jumping jacks, ten body-weight push-ups, ten T-Y-I shoulder motions, and ten chest-passes with a basketball.

#### Height

Height was assessed by having participants stand, fully erect and without shoes, next to a measuring tape on a wall. Participants stood with proper posture while the researcher recorded the height to the nearest 0.5 centimeter of the participant.

#### Body Mass

Researchers assessed body mass with a Detecto 437 eye-level physician’s scale. Participant’s body mass was recorded one at a time and in private. The participants stepped onto the scale while the researcher adjusted counterbalance weight to assess body mass. Body mass was measured to the nearest 0.25 kilogram.

#### BMI

Researchers calculated body mass index (BMI) using height and body mass. Body mass (kg) was divided by height (m) squared (15).

#### Seated medicine ball throw

The SMBT was conducted no longer than three minutes following the warmup protocol. To accomplish this, participants performed the warmup protocol and the SMBT in groups of five. After the researcher gave instructions on the warmup and SMBT protocols, participants performed the SMBT one at a time, in no particular order. Participants started by sitting at a 90° angle against a designated wall with their legs straight out and their head resting on the wall. Participants started by holding a 2 kg medicine ball against their chest. After receiving a verbal signal from the researcher, participants pushed the medicine ball in a chest-pass motion as forcefully as possible without their back or their head leaving the wall. In order to better identify the impact site of the ball, researchers lightly dusted medicine balls with gymnast chalk, which provided a mark on the floor where the ball initially made contact after the throw. The distance the medicine ball landed from the participant was then measured using a measuring tape. Prior to the throw, the measuring tape was placed on the ground, starting (0 meters) at the most distal point of the medicine ball when the participant completely flexed their arms (approximately 2 cm from the pelvis of the person performing the SMBT). The measuring tape recorded distance in increments of tenths of a meter from this point to the first point where the medicine ball landed. The measured distance was then recorded by hand using a data collection sheet. Each participant had three attempts to throw the medicine ball as far as possible with a two-minute break between each attempt. From the demonstration to the final attempt, the entire testing procedure took no longer than 45 minutes. This testing protocol is similar to that used in the studies by Margin et al. and Borms and Cools (4, 24). However, given the unique standardization of the current procedures, we refer to the current study methods as the “*Utah SMBT Protocol*”.

#### Design and Analysis

Researchers calculated means and standard deviations for distances thrown for age groups 12-13 and 14-15 and by gender. Researchers also calculated quartile rankings from mean distances to establish normative reference data. Pearson correlation coefficients (PCC or r) were calculated between trials for females and males. Likewise, PCCs were calculated for age and SMBT distance for each gender. Data was entered into Microsoft Excel and calculations made using said software.

## RESULTS

In total, 113 (56 males, 57 females) adolescents participated in the study. Tables 2 and 3 contain participant data including height, body mass, and BMI. Mean distances thrown by age group (12-13 and 14-15) and gender can be found in tables 4 and 5, for males and females, respectively. Quartile rankings by age and gender were calculated and shown in tables 4 and 5 for males and females, respectively. Mean scores increased with age groups (table 4 and 5), as did mean height and body mass (table 2 and 3).

**Table 2.** Male Participant Descriptive Information

| Age Group (years) | n | Height (cm) | Body Mass (kg) | BMI (kg/m<sup>2</sup>) |
| --- | --- | --- | --- | --- |
| 12-13 | 25 | 167.0±8.5 | 60.7±19.4 | 21.5±5.9 |
| 14-15 | 29 | 179.0±8.9 | 68.5±18.0 | 22.6±5.1 |

Note: mean ± standard deviation

**Table 3.** Female Participant Descriptive Information

| Age Group (years) | n | Height (cm) | Body Mass (kg) | BMI (kg/m<sup>2</sup>) |
| --- | --- | --- | --- | --- |
| 12-13 | 31 | 159.0±7.3 | 53.8±15.7 | 21.2±5.9 |
| 14-15 | 25 | 164.0±6.1 | 58.7±8.8 | 21.7±3.2 |

Note: mean ± standard deviation

**Table 4.** Male Mean Seated Medicine Ball Throw Scores and Quartile Rankings

| Age Group | Mean SMBT Score (m) | 25<sup>th</sup> Percentile | 50<sup>th</sup> Percentile | 75<sup>th</sup> Percentile | 100<sup>th</sup> Percentile |
| --- | --- | --- | --- | --- | --- |
| 12-13 (n = 25) | 4.3±0.7 | 3.9 | 4.2 | 5.0 | 6.6 |
| 14-15 (n = 29) | 5.2±0.8 | 4.3 | 5.0 | 5.8 | 6.8 |

Note: Percentile scores measured in the distance thrown in meters (m)

**Table 5.** Female Mean SMBT Scores and Quartile Rankings

| Group | Mean SMBT<br>Score (m) | 25<sup>th</sup><br>Percentile | 50<sup>th</sup><br>Percentile | 75<sup>th</sup><br>Percentile | 100<sup>th</sup><br>Percentile |
| --- | --- | --- | --- | --- | --- |
| 12-13 (n = 31) | 3.4±0.5 | 3.0 | 3.3 | 3.3 | 4.6 |
| 14-15 (n = 25) | 3.7±0.5 | 3.3 | 3.6 | 4.0 | 5.2 |

Note: Percentile scores measured in the distance thrown in meters (m)

Pearson correlation coefficients for between-trials comparisons for males and females ranged from r=0.85-0.97. Age significantly correlated with distance thrown in the SMBT (p=0.0001 r=0.455, p=0.0004 r=0.326 in males and females, respectively).

## DISCUSSION

The purpose of this study was to establish normative reference values for the SMBT. One hundred thirteen untrained male and female individuals aged 12-15 years participated in the study by throwing a 2 kg medicine ball with a 19.5 cm circumference. The study results included quartile rankings for the 12-13 and 14-15-year-old age groups in both males and females. Separating participants into age and gender categories was influential in establishing mean normative reference values. The establishment of quartile rankings can help guide further normative reference data research among this population.

In the current study, correlation coefficients for between-trials comparisons for males and females ranged from r=0.85-0.97 which ware similar to that noted by Beckham et al. (3). In previous research, Beckham et al. found a low magnitude of change (−0.02 to 0.08 m) between trial averages, a strong interclass reliability coefficient (ICC=0.97 - 0.99), and a low percentage of error for the SMBT (CV=3.2-3.9) when assessing twenty healthy undergraduate students using the SMBT with a ten-lb. medicine ball (3). These findings further suggest that the SMBT is a reliable measure of upper-body muscular power. Similarly, in a study by Borms et al., the SMBT showed strong test-retest reliability (r=0.980) in 29 male and female overhead athletes (age 21.6 ±2.5 years) using a two-kg medicine ball (5). Harris et al. found similar reliability in 33 older adults (age 72.4 ± 5.2 years) using a 1.5 kg ball (19). Davis et al. found that the test also yielded high reliability (r=0.88) in same-day trials and trials across two days in kindergarten-age children using a two-lb. medicine ball. (8).

All data in the current study was collected in a single day, as such, day to day reliability of the SMBT was not able to be determined. It is worth noting that the studies conducted by Beckham et al., Borms et al., Harris et al., Davis et al., as well as the current study, all positioned participants in a seated position with their back at a 90° angle (3, 5, 8, 19). This commonality suggests that positioning participants against a wall or flat surface will produce reliable results. While the mass of the medicine ball varies across studies, it appears that results will still show reliability provided that all participants use the same mass for all trials (3, 5, 8, 19).

The lack of standardized testing protocols acts as a limiting factor to the findings of most studies since the findings of each cited study are limited to only studies that share the same protocol. In addition to the lack of normative reference values, there is no official testing protocol for the SMBT. Some studies use protocols requiring participants to sit at a 45° on a bench (6, 10, 11, 20), while others require a 90° angle against a wall (4, 13, 24, 29).

When considering factors affecting maximum distance thrown using the SMBT, researchers should also consider chronological age. In the current study, age significantly correlated to distance thrown in the SMBT (p=0.0001, r=0.46 and p=0.0004, r=0.33 in males and females, respectively). As the age of participants in the current study approached maturity (19-25 years), throwing distance increased. The results of this study are consistent with the findings of previous research on the correlation between age and SMBT distance (1, 23). Researchers in a previous study recorded a significant (p<0.000) difference between male basketball players aged 11 and their 14-year-old peers in upper-body power on a laying medicine ball throw, further suggesting a correlation between age and throwing distance (1). In the case of the basketball players, throwing distance increased with age. As players’ ages approached maturity (19-25 years), throwing distances increased (1). The findings of previous studies have found that as participant age moves away from age 25 in either direction, throwing distance decreases (1, 9, 23).

Gender is another consideration when assessing muscular power. The results of this study are consistent with previous research in terms of the effect of gender on SMBT distance. Participants in the male group of the present study scored significantly (p=0.009) higher than the female group. In a previous study, Lockie et al. found that female recruits of a law enforcement agency performed lower on the SMBT than their male counterparts (p<0.001) (23). British boys (age 4-7) scored significantly higher on the SMBT than girls in the same age group (12). Similarly, a study by Hacket et al. suggested that the SMBT is a stronger predictor of muscular power when comparing results to participants of the same gender (18). Normative reference values for the studies mentioned above were either not calculated or not reported, thereby limiting direct comparisons (12, 18, 23, 28).

Limitations for this study include the participant sample sizes and characteristics, lack of geographical diversity, and the assumption that all participants gave maximal effort. The small sample size may have increased standard deviations of scores and raises questions of external validity. Additionally, all participants were from the same school within the state of Utah, United States (i.e. *Utah SMBT Protocol*). The participants in the study were 95% white with various other ethnicities represented in the remaining 5%. Considering the lack of diversity of the population, it is possible that a more diverse population would affect results of future studies. The ages of the participants were 12-15 years, meaning that the norms established will only apply to those age groups in males and females, respectively. It is assumed that all participants were untrained in the present study but resistance-training status may have varied between individuals and groups.

While the current study assumes that all participants gave maximal effort for every attempt, there is no metric to prove that assumption. Researchers instructed participants to use maximal effort for every throw, however the inability to quantify whether participants gave maximal effort could limit the reproducibility of data. Encouraging participants to give maximal effort for every attempt will improve validity and reliability of results in future studies, however similar limitations will persist. In the future, a detailed reliability analysis of the data collected in the present study utilizing the Utah SMBT Protocol should be undertaken, as did Beckam et. al. (3).

Perhaps the single most significant limiting factor for this study was the COVID-19 virus. This study used a single school location to limit contact between individuals and help stop the spread of the COVID-19 virus. Precautions were required to implement effective social distancing, sanitizing, and limited exposure. These precautions included limiting how many locations the researcher(s) traveled to, however utilizing multiple locations would have likely increased the sample size and positively impacted the robustness of the data. These precautions and several others limited the number of individuals that could participate and the final sample size.

Future research should validate or adjust the quartile rankings for the population used in this study. The same protocol and medicine ball must be used to reproduce or validate the findings of this study. This study used a 2 kg medicine ball with a 19.5 cm diameter, and participants sat at 90° during the Utah SMBT Protocol. Future research should aim to gather a larger sample size and complete the same procedures to validate and expand on the reference norms. In addition, normative reference values might particularly be valuable in high-school-age individuals. This information would provide baseline metrics by which coaches and educators could compare levels of either trained athletes or untrained individuals. This information could be used to facilitate better training for upper-body muscular power gains. Coaches and educators would also improve ability to assess readiness for sport at the high school level.

## CONCLUSION

This study has produced an initial set of normative reference values for male and female adolescents aged 12-15 for the Utah SMBT Protocol. Males age 12-13 had a mean score of 4.3±0.7 m, while males age 14-15 had a mean score of 5.2±0.8 m. Female participants age 12-13 had a mean score of 3.4±0.5 m, and females age 14-15 threw for a mean score of 3.7±0.5 m. This normative reference data was established with participants seated at 90° and using a 2 kg medicine ball with a 19.5 m diameter.

## APPLICATIONS IN SPORT

This research supports the use of the Utah SMBT Protocol as a means for coaches, athletes, and strength and conditioning professionals to assess the upper-body muscular power of adolescent individuals in a safe, effective, and efficient manner. Researchers can use this test as a baseline and formative assessment to measure upper-body muscular power in adolescents. This research also helps to establish procedures for further normative reference data gathering. It is important to note that replication of the test used in this study would require participants to sit at 90° and use a 2 kg medicine ball with a 19.5 cm diameter.

## ACKNOWLEDGMENTS

We would like to thank the student participants, parents, and school district administrators for their dedication to sport science and contributions to this project.

---

**Figure 1.**
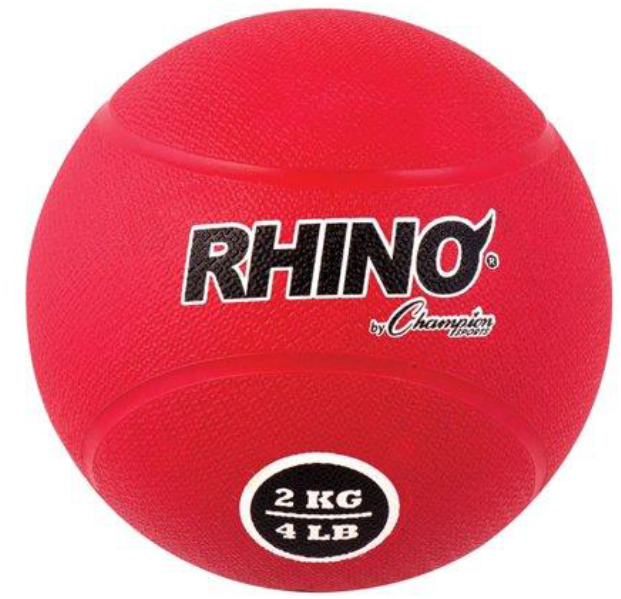
Example of medicine ball from https://www.walmart.com/ip/Champion-Sports-2kg-Rubber-Medicine-Ball/42844198

